# Machine learning applied to otolith microchemical data to discriminate stock of origin in salmon

**DOI:** 10.64898/2025.12.01.691230

**Authors:** Ben Makhlouf, Daniel E. Schindler, Valentina Staneva

## Abstract

A key challenge in ecosystem-based fisheries management is sustainably harvesting co-occurring species and populations that differ in their vulnerability to fishing. This challenge is exemplified in western Alaska Chinook salmon, where recent population declines have led to closures of in-river subsistence fisheries, but identifying the cause of these declines is limited by an inability to identify the river of origin for marine-caught fish in offshore fisheries that target more abundant species and populations. This problem is particularly acute for estimating the impacts of bycatch on individual salmon populations which have demographic independence but no genetic differentiation from which stock assignments can be made. Here we use machine learning approaches to assess the efficacy of using the microchemical history preserved in fish otoliths to assign individuals to their river of origin in western Alaska. We tested the classification ability of three machine learning algorithms (Random Forest, K-Nearest Neighbors, and Support Vector Machines) combined with two time series smoothing techniques (Moving Average and Generalized Additive Models) to classify Chinook salmon to their river of origin using otolith time series data. Model accuracy ranged from 71.9% to 92.5%, with optimal performance achieved by Random Forest applied to GAM-smoothed data. Watershed-specific performance ranged from 86.8% to 93.7%, with most misclassifications occurring between spatially proximate Kuskokwim and Nushagak rivers. Raw predicted probabilities from classification algorithms were calibrated to reflect true class probabilities, enabling the incorporation of model results into decision analyses with explicit consideration of misclassification risk tolerance. The success of these models offers immediate utility for estimating marine mortality impacts across the region’s three major river systems as well as an opportunity to understand commercial fisheries impacts on individual populations at substantially finer spatial scales than had been previously possible.

## Introduction

In remote, large, and complex ecosystems, monitoring ecosystem processes using conventional methods can be limited. This, in turn, is a serious impediment towards developing ecosystem-based approaches to management that accounts for variation in the vulnerability of individual species and populations to human stressors. Management of Pacific salmon in western Alaska exemplifies this challenge. Adult salmon in the Yukon, Kuskokwim, and Nushagak river-basins migrate through remote, complex, and rugged ecosystems. Each of these river-specific populations contribute towards overall regional population health, however, population dynamics among them vary both within and among return seasons. Individuals from these populations comingle in the ocean where they are intercepted by marine fisheries targeting other, more abundant species (DeFilippo et al., 2025; Stram & Ianelli, 2015). However, the potential impact on ecosystem health depends on the relationship between marine mortality and river-specific population dynamics. As such, monitoring at the basin or sub-basin scale is important towards assessing the health of river-specific stocks as well as their vulnerability to marine fisheries.

Salmon populations in western Alaska have historically supported valuable commercial fisheries and have served as an important subsistence resource for upstream communities. However, recent declines in the number of Chinook salmon (*Oncorhynchus tshawytscha*) and chum salmon (Oncorhynchus keta) returning to western Alaska watersheds have led to fishery restrictions and a crisis of food insecurity and cultural loss (DeMaster et al., 2025; Feddern et al., 2023). Several marine-based hypotheses have been proposed for declining population health including mortality due to changing environmental conditions or bycatch in federal groundfish fisheries and mixed stock salmon fisheries in Alaska waters (Lamborn et al., 2025). Evaluating the impact of these hypotheses requires evaluating marine mortality against river-specific population dynamics. However, genetic stock identification (GSI), a powerful and widely used tool for estimating the provenance of Chinook salmon stocks for in-season management (Beacham et al., 2004; Hess et al., 2014), cannot discriminate among watersheds in many cases (Larson et al., 2014; Templin et al., 2011; Van Doornik et al., 2024). Although fish from the middle and upper portions of the Yukon can be reliability distinguished using GSI, the Coastal Western Alaska (CWAK) genetic reporting group contains the entire Kuskokwim and Nushagak river-basins as well as a large portion of the Yukon (Figure 1a). As a result, the watershed of origin cannot be reliably distinguished for fish assigned to this genetic baseline, hampering the ability to quantify the impact of marine-based fishing mortality on river-specific population dynamics.

**Figure 1:**
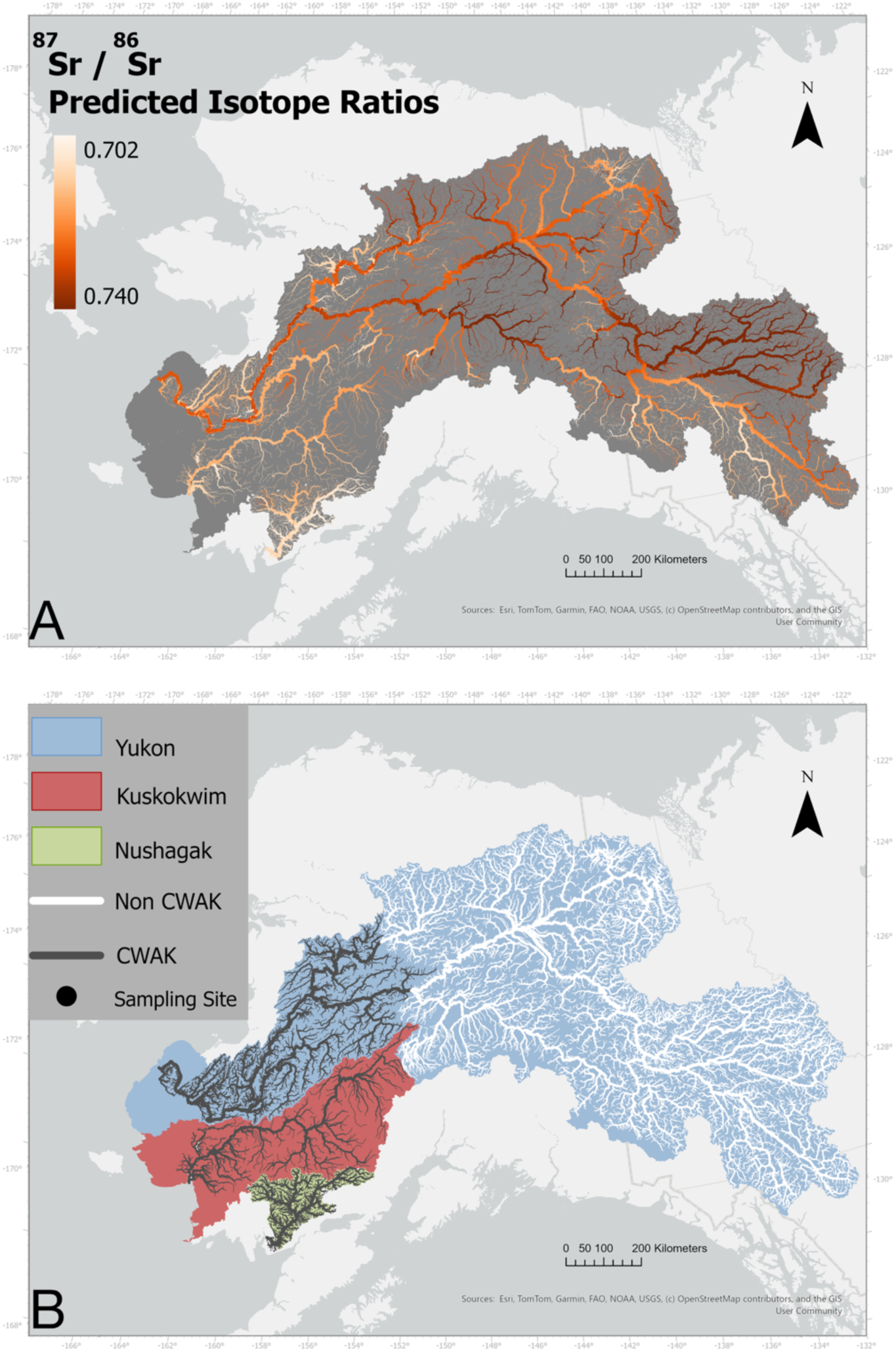
**A)** Three major river systems in Western Alaska, the Yukon (Blue), Kuskokwim (Red), and Nushagak (Green) watersheds. Black circles indicate sampling locations for otolith collection (and genetic collection in the Yukon) on each river. **B)** ⁸⁷Sr/⁸⁶Sr isoscape of all three river-basins. Values range from 0.702 to 0.740, with darker colors indicate more enriched values whereas lighter colors indicate less enriched.

Isotope-based methods for estimating provenance using ear stones found in fish, or otoliths, have emerged as an alternative approach to GSI, offering finer spatial resolution than genetics in some cases (Brennan et al., 2015; Brennan & Schindler, 2017; Kennedy et al., 1997). The relative ratio of strontium-87 to strontium-86 (⁸⁷Sr/⁸⁶Sr) is heterogeneously distributed across geologically diverse landscapes, faithfully incorporated into the metabolically inert otolith structure, and has been demonstrated to be temporally stable in Alaskan watersheds (Bataille & Bowen, 2012; Brennan et al., 2015; Campana, 1999; Capo et al., 1998). As such, this method has been used to estimate provenance of western Alaska salmon from within the watersheds where they are harvested by comparing the ⁸⁷Sr/⁸⁶Sr value isolated from the natal freshwater region of the otolith to strontium values modeled across the landscape (Brennan et al., 2015, 2019) However, this method is limited by the level of isotopic heterogeneity relative to the spatial scale of interest (Makhlouf et al., 2025). In western Alaska, the heterogeneity in strontium isotope ratios is sufficient to discriminate among major tributaries within a single watershed from which an individual fish is captured, however, redundancy in strontium isotope ratios among rivers at the regional scale prevents reliable assignments of river of origin (Figure 1b.). As a result, determining the river of origin from a single ⁸⁷Sr/⁸⁶Sr value is distinctly limited for marine caught fish which could have originated from any of the three watersheds.

Otolith growth and isotopic incorporation occur continuously over an individual’s lifetime (Campana, 1999). When analyzed along the growth axis of the otolith, this results in a ⁸⁷Sr/⁸⁶Sr time series that encodes maternal migration history, sequential habitat use, and relative growth of the individual as it migrates through the river (Hegg et al., 2019; Reis-Santos et al., 2022). In some cases, however, rapid movement through multiple environments prevents full isotopic equilibration between the otolith and ambient water. For Pacific salmon, which often migrate quickly among different freshwater rearing habitats on their migration to the ocean, otolith strontium data following the natal origin region may not reflect the true ⁸⁷Sr/⁸⁶Sr values of each outmigration habitat. Instead, these data provide a series of transient patterns produced by incomplete and ongoing turnover dynamics within each habitat. The shape of these patterns, and the relationships among them, may encode distinctive river-specific life histories that together form a unique “fingerprint” of river-specific stocks. Because these patterns are subtle and the number of possible combinations in large river basins is high, such identifiers are difficult to detect using traditional statistical or manual approaches. Machine learning (ML) methods, which are particularly well-suited to interpreting complex, multidimensional patterns in data, have only recently been applied to otolith chemical analysis. However, their limited applications have shown promising performance relative to traditional techniques (Arai et al., 2023; Mercier et al., 2011). At the same time, there is increasing recognition that reducing continuous otolith laser ablation (LA) ⁸⁷Sr/⁸⁶Sr data to a single value discards potentially powerful contextual information about life history strategies (Arai et al., 2025 (in review); Hegg & Kennedy, 2021). Combining the temporal resolution of otolith time series data with the statistical power of ML offers the potential to detect otherwise indiscernible patterns within and across LA datasets. With the addition of known watershed origin labels, these methods could be used to develop a model capable of discriminating the river of origin for Chinook salmon from western Alaska caught in marine fisheries.

Here, we apply ML classification algorithms to a large dataset of otolith LA data from Chinook salmon in western Alaska with a known river of origin. In doing so, we test the potential of ML methods to identify watershed specific characteristics to discriminate among major rivers and develop a model to reliably estimate river of origin for marine-caught Chinook salmon in western Alaska.

## Methods

### Otolith and genetic data

Chinook salmon otoliths were collected from the Yukon (2015, 2016, 2019–2021), Kuskokwim (2017–2022), and Nushagak (2011, 2014, 2015) river basins at test fisheries near the downstream terminus of each river (Figure 1). Fish were sampled through collaborative efforts with the Alaska Department of Fish and Game Lower Yukon Test Fishery (LYTF), Orutsararmiut Native Council at the Bethel Test Fishery, and from Chinook salmon caught within the Nushagak commercial fishing district in Bristol Bay (Black dots, Figure 1). We assumed that all Chinook salmon caught in this fishing district were returning to the Nushagak River. For each year approximately 500 otoliths were collected throughout the season. From these, ∼250 otoliths from each year were selected for further analysis to ensure equal representation relative to CPUE across the full run.

Selected samples were cut along the transverse plane using a low speed isomet saw to reveal the growth structure of the otolith, mounted on microscope slides, and polished to enhance clarity (Donohoe & Zimmerman, 2010; Zimmerman & Reeves, 2002). Samples were then analyzed using Laser Ablation Inductively Coupled Plasma Mass Spectrometry (LA-ICP-MS) at the University of Utah Strontium Metals Lab, with the exception of 2022 Kuskokwim samples which were processed at the W.M. Keck Collaboratory for Plasma Spectrometry at Oregon State University. Laser ablation was performed from the otolith core to just after the onset of the marine signature along the dorsal lobe scanning at 2 microns/second. The resulting dataset includes a concurrently collected time series of ⁸⁷Sr/⁸⁶Sr and Sr^88^ along the same transect for all individuals. Because the Yukon river-basin contains multiple distinct genetic reporting groups, fin clip samples were collected from a subset of Yukon individuals for genetic stock assignment. From these samples, a posterior probability of occupancy for three genetic reporting groups (Lower, Middle, and Upper Yukon) was assigned following methods outlined in Makhlouf et al., 2025 (West & Dann, 2019). Each individual with available genetic data was then assigned to the reporting group with the highest probability of origin.

### Data pre-processing

To standardize all data to the same life history period full laser ablation datasets were trimmed to include only data between the onset of the natal origin region and the beginning of marine residence. These cutoff points were identified by examining patterns in ⁸⁷Sr/⁸⁶Sr, ⁸⁸Sr, and visually apparent morphological features along the laser ablation line (Brennan & Schindler, 2017). The natal origin region was manually identified by the first steady state after the decline in ^88^Sr and movement of ⁸⁷Sr/⁸⁶Sr away from the global marine value (0.7092), which represents the maternal signal in the otolith. Although we hypothesized that the maternal signature would differ among watersheds, and thus could be useful for classification, the start of the laser ablation transect and quality of the core signal varied among samples and years. Therefore, to maximize consistency among years we did not include the portion of the time series before the natal origin region. A cutoff at the marine entry was identified by the marine check on the otolith (dark annuli region), a rapid increase in ⁸⁸Sr concentration, and a shift in ⁸⁷Sr/⁸⁶Sr towards the global marine value at the end of the time series. Manual selection of trimming locations was randomized and blinded to avoid bias. All trimmed datasets were then manually checked for quality and samples with significant error (i.e large cracks) or without a clear core and marine signal were removed from the dataset. The resulting dataset contained time series from 2,531 individual otoliths (971 Yukon, 875 Kuskokwim, and 685 Nushagak).

To test whether time-series smoothing techniques could improve classification model performance, we applied two commonly used approaches: moving average (MA) and a generalized additive model (GAM) on the raw strontium isotope data. For the moving average smoothing we used a 60-point center weighted window. The GAM was fitted using thin plate regression and penalized iteratively re-weighted least squares (P-IRLS) to maximize goodness of fit, with a generalized cross-validation (GCV) method used to minimize overfitting (Wood, 2006). The maximum degrees of freedom (k) were scaled by the number of data points in each ablation line (n) by k = 10 * n ^2/9^ (Kim and Gu, 2004, Brennan et al., 2014). This process resulted in three versions of the same laser ablation time series for each individual: GAM-smoothed, MA-smoothed, and raw LA data. Finally, all samples were interpolated to a common length defined as the average length of all time series of that type (694 points for non-MA and 664 points for MA-smoothed samples, respectively) (Hegg and Kennedy, 2021). This was done to satisfy the requirement for equal length vectors for ML algorithms while minimizing warping from interpolation.

Principal component analysis (PCA) was applied to all timeseries from the dataset in the top-performing model (GAM smoothed data) to qualitatively assess clustering patterns in multivariate space (Figure 2). PCA transforms the original time series into uncorrelated principal components that capture the greatest variance in the dataset, thereby highlighting variability among time series that may contain distinguishing watershed characteristics. To further illustrate how this variability among time series can distinguish among individuals that are otherwise indistinguishable using the natal freshwater ⁸⁷Sr/⁸⁶Sr, PCA analysis was also done on a subset of the GAM smoothed dataset which include only individuals with a natal ⁸⁷Sr/⁸⁶Sr between 0.7080 and 0.7085. PCA plots were produced to visualize clustering of both PC1 (47.5% of variance) and PC2 (20.9% of variance) as well as between PC1 and PC3 (10.2% of variance) (Figure 3a). We then displayed the absolute values of each point’s loading on the corresponding index of the time series to identify regions of the time series which contribute to interclass separation among individuals with the same natal origin ratio (Figure 3b).

**Figure 2:**
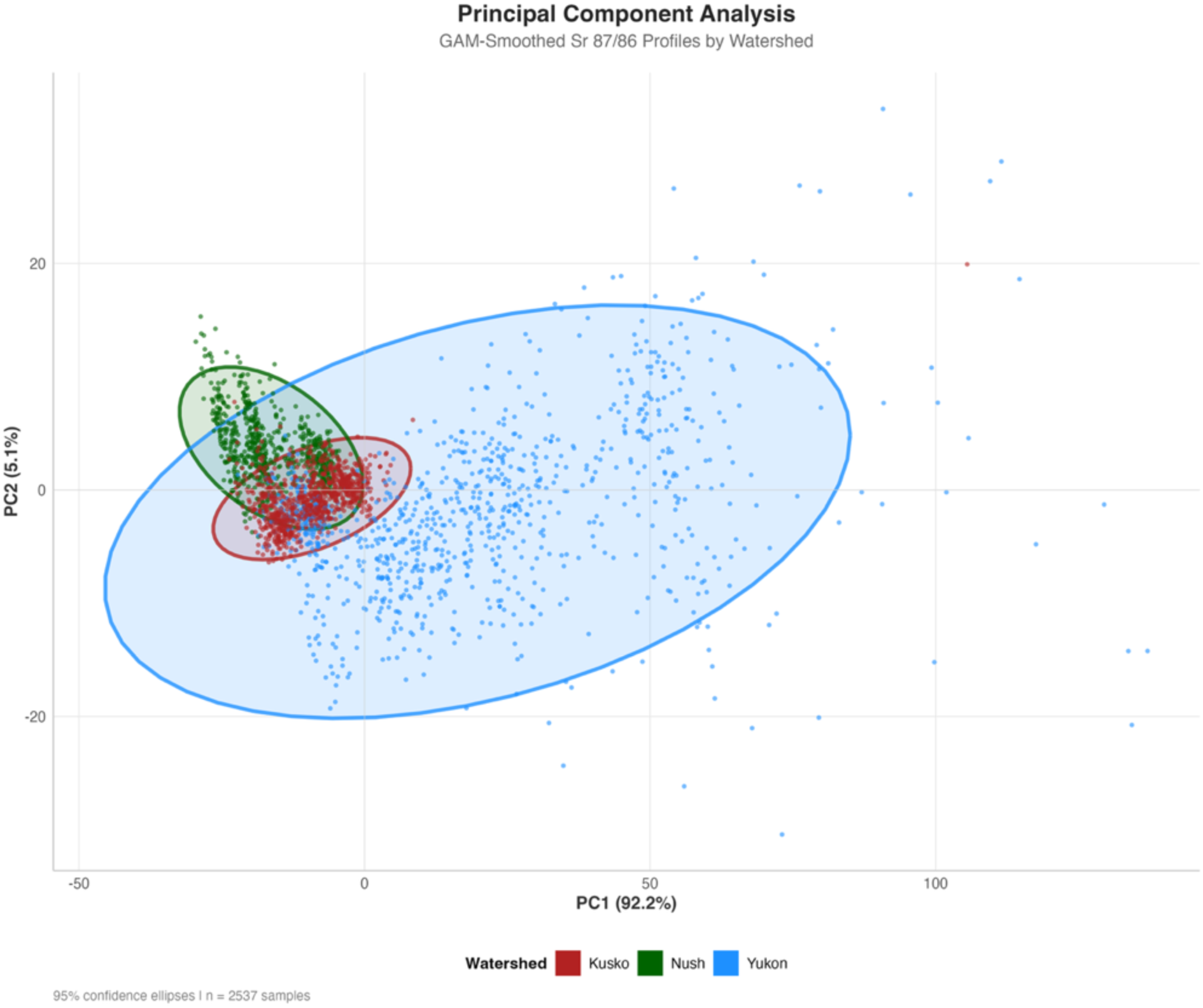
Principal Component analysis of GAM smoothed timeseries data (n = 2,531 otoliths). Significant clustering is evident, but with overlap between all watersheds. Yukon (Blue points) contain a much larger range of values, indicative of the large range in natal ⁸⁷Sr/⁸⁶Sr values in the basin. Kuskokwim (Red points) overlap between both Nushagak and Yukon fish, likely due to its position in the region between both watersheds.

**Figure 3:**
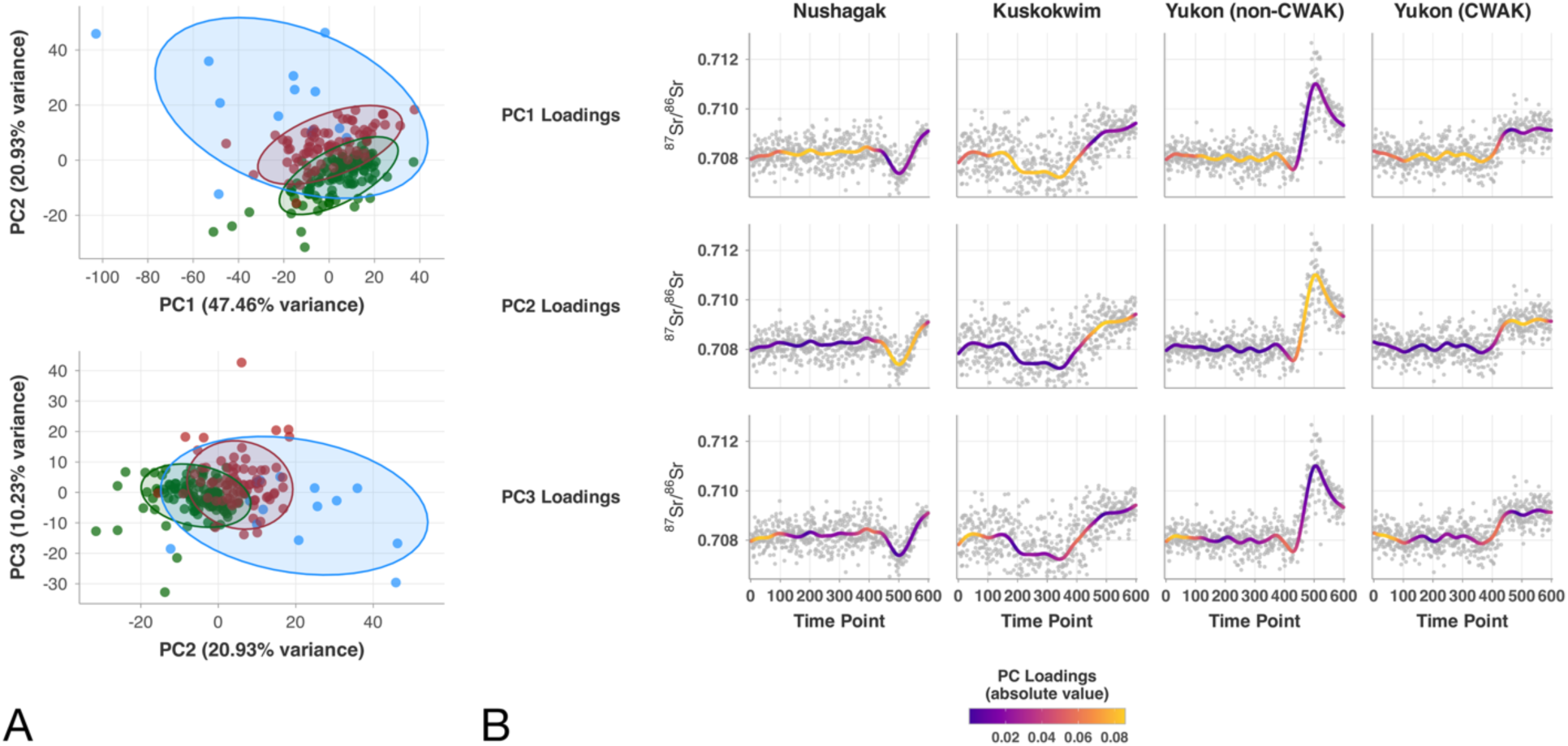
A) Principal component analysis of all samples with a natal origin ⁸⁷Sr/⁸⁶Sr of 0.7080 - 0.7085. Clustering among watersheds is evident between principal components 1 and 2 (68.39% of variance), as well as between principal components 2 and 3 (31.16% of variance). There is some overlap between all three watersheds in both cases. B) Sample otolith LA timeseries colored by the absolute value of the PCA loadings at associated timeseries indices. PC1 (top row) shows high loading on the middle section of the timeseries after the natal origin region. PC2 is driven by the region of the timeseries towards the marine entry, likely the result of mainstem migration to the ocean. Separation along PC3 is driven by the very beginning of the timeseries, which is the region associated with the natal origin. Notably, Yukon individuals from the CWAK region contain a less pronounces mainstem signal as compared to their counterpart from the upper or middle Yukon genetic groups.

### Machine Learning Classification Models

All individuals in the dataset were randomly partitioned into either training or testing datasets using a randomized 80/20 split. To ensure consistency across analyses, the same individuals were assigned to the training and testing sets for all three datasets (non-smoothed (Raw), MA, and GAM). We trained three supervised classification algorithms: K-Nearest Neighbors (KNN), Random Forest (RF), and Support Vector Machine (SVM). The Random Forest model was configured with 500 trees, and the number of predictors randomly sampled at each split (mtry) was set to the square root of the total number of predictors, following standard practice for classification problems. The KNN model was trained using five neighbors, while the SVM model used default hyperparameters for the RBF kernel. All models were implemented using the tidy models framework in R (Kuhn & Wickam, 2020).

Trained models were then evaluated on two separate versions of the testing dataset for each data type. First, models were tested on the original randomly selected 20% holdout set. Second, models were evaluated on a modified testing set in which samples with natal origin ⁸⁷Sr/⁸⁶Sr values above 0.7130 were excluded. Only the Yukon River watershed contains enriched isotope values above this threshold and samples with a natal origin in this range can therefore be distinguished using natal origin values alone. This modified dataset, hereby referred to as the “Restricted” testing set evaluates the ability of the model to correctly classify only samples that are otherwise indistinguishable using natal origin values alone. Model performance was assessed using accuracy and F1-score (Figure 4). Accuracy measures the proportion of correctly classified samples:

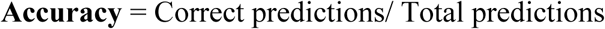

The F1-score balances precision and recall and is calculated as their harmonic mean:

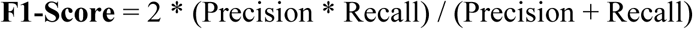

**Figure 4:**
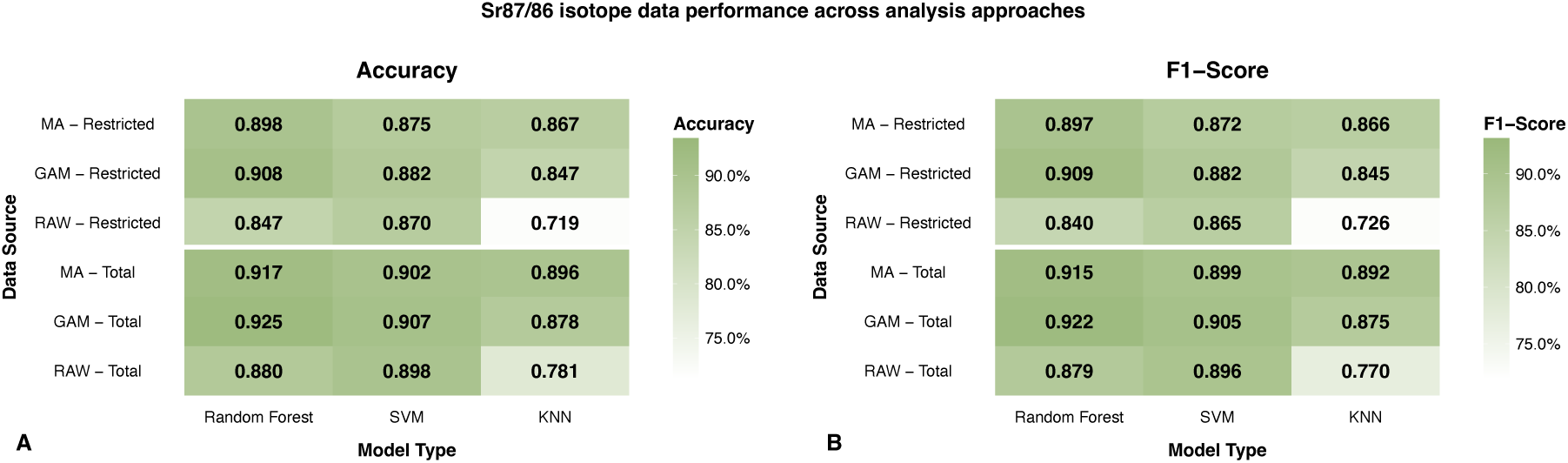
Model Performance comparison among data preprocessing steps and model types. Accuracy indicates the proportion of each class correctly classified. F1-Score is the harmonic mean of precision and recall, providing a balanced performance measure across all watershed classes. Results compare two ⁸⁷Sr/⁸⁶Sr timeseries smoothing methods (GAM, MA) and three machine learning algorithms (Random Forest, SVM, KNN) using either all test data (Total) or only overlapping isotope ranges (Restricted). Darker green colors indicate higher scores.

Here, precision is the proportion of samples predicted as positive that are truly positive, and recall is the proportion of actual positive samples that are correctly identified. The F1-score provides a single metric that accounts for both false positives and false negatives, making it particularly useful for evaluating performance on imbalanced datasets (Van Rijsbergen, 1975). For the highest-performing model, confusion matrices were generated to visualize class-specific accuracy and to explore trends in misclassification among classes (Figure 5). In addition to calculating the accuracy by watershed, we calculated the classification accuracy of fish from within the CWAK (Coastal Western Alaska) genetic reporting group by removing individuals from the testing set with genetic assignments to the Middle or Upper Yukon reporting groups, which are outside of the CWAK region.

**Figure 5:**
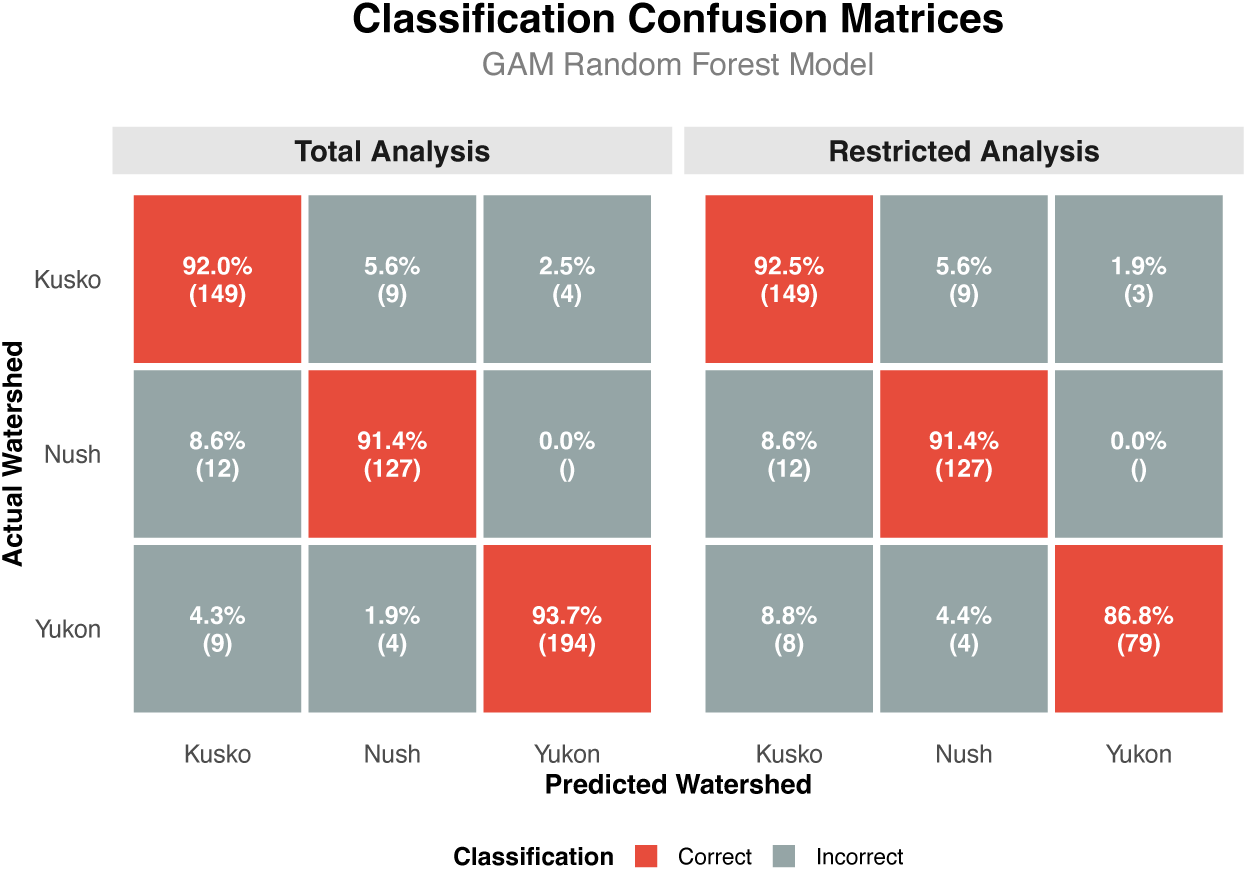
Confusion matrix from the highest performing model (Random Forest on GAM Smoothed data) for both total test set and overlapping subset. Boxes along the horizontal axis indicate correctly classified individuals (red). Vertical axis labels are the actual watershed labels whereas horizontal labels are the predicted class from the model. Percentages indicate the percent of the actual class identified as the predicted class and numbers indicate the number of the actual class assigned to the predicted class.

### Probability calibration and threshold testing

Many ML classification models produce predicted probability estimates (confidence scores) that correctly identify rank class likelihood but produce estimated probability values that do not accurately reflect the observed probability of correct prediction (Dormann, 2020; Silva Filho et al., 2023). To be treated as reliable confidence values, we should expect that the observed proportion of individuals belonging to a given class aligns with the model’s assigned probability for that class. For example, if our model predicts individuals to have originated from the Kuskokwim watershed with 80% probability, then approximately 80% of those individuals should in fact be Kuskokwim fish. However, this assumption is oftentimes not met, limiting the utility of these values for decision-making which relies on confidence thresholds of correct assignment probability or propagation of uncertainty through further probability-based models. To calibrate predictions from our models, we used the {probably} package in R to apply multinomial isotonic regression to our testing set predictions (Kuhn et al., 2025). This approach adjusts the predicted probability scores to better align with the true observed probabilities through a non-parametric monotonic transformation. We verified calibration effectiveness through a visual assessment of plots showing the predicted probabilities against observed class frequencies across probability bins (ex. Supp. Fig 1.). Calibration transformations were fitted separately for each model-data type combination ensuring that probability estimates from our models accurately reflect observed classification probability. Using the calibrated testing set probabilities, we then tested the classification accuracy obtained from our best model above calibrated confidence thresholds ranging from 60% to 90%. For a given confidence threshold, we calculated the number of individuals assigned to a the correct class with a calibrated probability value at or above the given threshold. This value was then compared to the total number of individual mis-identified as another class or assigned to the correct class but with a calibrated probability value below the threshold. Class-specific and overall accuracies were recorded for each threshold value (Figure 6).

**Figure 6:**
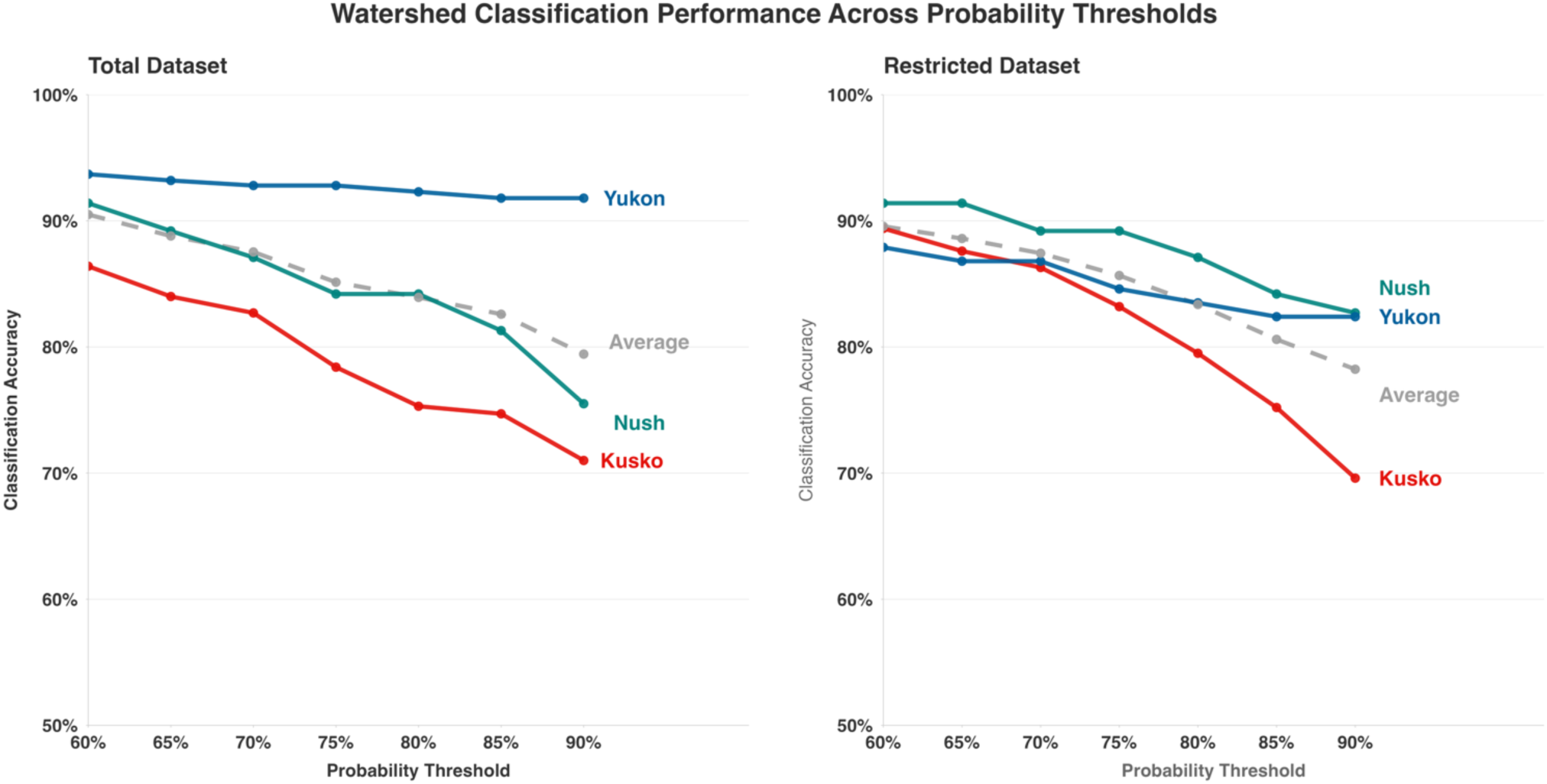
Classification accuracy with different probability thresholds (60-90%). Points indicate the percentage of correctly classified individuals with a calibrated probability value above the given threshold value. Grey dashed lines represent the overall accuracy, whereas colored lines illustrate the class-specific probability at that threshold

## Results

### Principal Component Analysis

Principal component analysis of the full GAM-smoothed dataset (n = 2,531) revealed significant clustering by watershed. The first two principal components together captured 97.3% of the total variance in the dataset (PC1: 92.2%, PC2: 5.1%, Figure 2). Despite clear clustering patterns, some overlap remained among all three watersheds. Kuskokwim samples (Figure 2, red) exhibited the greatest degree of overlap with both Nushagak and Yukon samples. Yukon samples (Figure 2, blue) covered a much larger region of the principal component space, reflecting the broader range of potential isotope values that create unique time series compared to the more constrained ranges observed in the other two watershed classes

The pattern observed in the full dataset was also found among only individuals with the same natal origin value (0.7080 – 0.7085) (Figure 3a). The PCA results for this subset displayed similar clustering results, with a larger overall range of Yukon samples and overlap between all three watersheds. Mapping the loadings of each principal component onto the time series revealed distinct regions driving class separation. The first principal component showed high loading values in the middle portion of the time series, potentially encompassing both the end of the natal origin region as well as secondary rearing locations. The second principal component was driven by variation occurring just before the onset of the marine region, likely representing residency in mainstem tributaries during oceanward migration. The relative shape at this point in the time series likely reflects differences in mainstem tributary ⁸⁷Sr/⁸⁶Sr values among the three watersheds ( 0.714, 0.709, 0.707 in the Yukon, Kuskokwim, and Nushagak, respectively). In many samples, this difference in mainstem value appears to have resulted in either a dip in ⁸⁷Sr/⁸⁶Sr for Nushagak fish, an increase for Yukon fish, or a more subtle increase for Kuskokwim fish before the marine cutoff (Figure 3b). Notably, many Yukon fish assigned to the CWAK region appeared to have a much less pronounced increase in ⁸⁷Sr/⁸⁶Sr, which is consistent with less residence time in the mainstem as a result of a much shorter distance from their natal origin region to the ocean. Principal component 3 appears to be driven by values at the very beginning of the time series corresponding to the start of the natal origin signature as well as some variation in values just before the mainstem region described above. There is minimal separation by watershed along this principal component (Figure 3a), which is expected given that all individuals in this subset have very similar natal origin values.

### Model Performance

Assignment accuracy across both testing sets ranged from 71.9% to 92.5% and F1-Score values ranged from 72.6% to 92.2%. The highest accuracy and F1-score was achieved by the random forest model applied to the full GAM smoothed testing set (Figure 4). Smoothed time series data outperformed the raw timeseries in all cases, indicating that the noise in the raw data may have inhibited model performance by obscuring meaningful patterns. Among model types KNN consistently performed the lowest in both accuracy and F1-Score. In general, RF analysis outperformed both KNN and SVM, however, SVM was the highest performer on raw datasets (89.8% accuracy on total testing set and 87.0% on restricted testing set. Unsurprisingly, the restricted testing set performance was slightly lower than that of the total training set, with the highest classification accuracy of 90.8% as compared to 92.5%. These results are expected given the relatively straightforward discrimination of highly enriched Yukon samples which were removed in the restricted comparison.

The best performing model (RF, GAM-smoothed data) displayed some variation in overall accuracy among classes. In the total testing set 92.0% of Kuskokwim, 91.4% of Nushagak, and 93.7% of Yukon samples were correctly identified (Figure 5). By percentage of each class, the most common misclassification was Nushagak fish classified as Kuskokwim (8.6% of Kuskokwim fish) followed by the inverse (5.6% of Nushagak fish). Interestingly, after removing the most discernable Yukon fish (⁸⁷Sr/⁸⁶Sr > 0.713), Yukon classification accuracy dropped to 86.8%. Classification accuracy for Nushagak fish did not change (91.4%) whereas Kuskokwim saw a slight increase to 92.5%. In this testing dataset (restricted) the most frequent misclassification was Yukon fish predicted to have been Kuskokwim (8.8% of Yukon fish), however this represented only one additional individual misclassified (9 in the total analysis vs. 8 in the restricted analysis). Classification accuracy within the CWAK reporting group, which includes the Kuskokwim, Nushagak, and Lower Yukon regions was 90.0%. Classification accuracy for regions in our dataset not within CWAK was 97.9%. Yukon classification accuracy for those within the CWAK group fell to 73.3%. However, it should be noted that the sample size of CWAK Yukon individuals was significantly lower than other groups ( n = 29 ), resulting in a more pronounced impact to class-specific accuracy from misassignment to relatively few individuals.

### Probability tuning and threshold analysis

Probability tuning was used to align predicted probabilities from classification algorithms with observed probabilities by calibrating model outputs to better reflect true class likelihoods. Visual assessments indicated improved calibration, with the relationship between predicted and observed probabilities approximating a 1:1 line across all classes and model–data type combinations (ex. Supp. Fig 1). Notably, many of the predicted probability values from our models appeared to follow this assumption relatively well even before calibration. Across the range of tested thresholds, overall classification accuracy ranged from 91 – 79% for the total testing set and from 89.5% to 78% for the restricted dataset after calibration (Figure 6). This reflect the declining number of individuals which can be assigned be correct defined by the model with an added requirement of a high probability value. In the total testing dataset, Yukon classification accuracy remained well above 90%, even at the very high (>90%) confidence threshold. Nushagak and Kuskokwim classification accuracy, however, fell steadily as the threshold was raised. Kuskokwim classification accuracy dropped steadily from 87% to 71% whereas Nushagak accuracy declined more quickly at higher probability thresholds from 92% to 75%. The restricted dataset contained slightly different trends, with Yukon accuracy remaining relatively stable between 89% and 83% over the full range of threshold values. In this testing set, Nushagak accuracy dropped steadily from 91% to 84%, whereas Kuskokwim accuracy declined rapidly from 89% to 69%.

## Discussion

Machine learning tools applied to otolith laser ablation time series data proved highly effective for classifying western Alaska Chinook salmon to their rivers of origin when trained on a large dataset of known-origin individuals. We selected three methods (KNN, RF, and SVM) to represent a spectrum of commonly used ML techniques, ranging from the relatively simple KNN to more complex, nonlinear approaches like RF and SVM. Among these, random forest algorithms emerged as particularly powerful and robust, achieving over 90% classification accuracy across the smoothed datasets. Notably, even our simplest model (KNN) performed well, delivering reasonably high classification accuracy across both testing sets. In addition, datasets preprocessed with time series smoothing techniques significantly outperformed raw laser ablation data. This is unsurprising given the inherent noise in otolith LA data, however, these results demonstrate the added value of applying time series preprocessing techniques to otolith LA datasets and suggests that even greater discriminatory power may be possible using additional, more complicated approaches. Given the relatively small sample size in our testing set, we acknowledge that the relatively subtle difference in classification accuracy among methods may not be significant. However, even if the difference is insignificant, all methods presented here provide a high level of classification accuracy.

Using the best preforming model, watershed-specific classification accuracies on the full testing dataset exceeded 90% for all three watersheds. Most misclassifications occurred between Nushagak and Kuskokwim fish, consistent with expectations given the greater overlap in ⁸⁷Sr/⁸⁶Sr values in potential strontium sources and closer spatial proximity between these watersheds. This limitation is further illustrated in the confidence threshold analysis, in which we imposed the additional requirement that the model must assign to the correct class with a calibrated probability value above a given threshold. Preforming this analysis on the total dataset resulted in a significant decrease in classification accuracy in the Nushagak and Kuskokwim as we increased the threshold probability value. This reflects the fact although the model correctly assigns individuals to these watersheds in most cases, the associated confidence of the model is much lower for these classes given their similarity in time series shape. In contrast, Yukon fish exhibited unique patterns in otolith LA data due to the watershed’s extreme spatial scale, broader range of isotope values, and extended migration timing (Figure 2). This likely resulted in otolith time series which were distinct compared to the other watersheds either because of natal origin values only found in the Yukon (⁸⁷Sr/⁸⁶Sr > 0.713) or prominent features resulting from a longer migration through the enriched mainstem tributary (i.e Figure 3b, column 3). Yukon classification accuracy dropped substantially in the restricted testing set (86.86% vs. 93.7% in the total testing set), despite similar numbers of misidentified individuals. This decline likely reflects the removal of the most easily classified cases, which is further supported by higher misclassification rates among CWAK individuals compared to those from the middle and upper Yukon regions. These challenges likely stem from shorter migration patterns that include less residence time in the mainstem Yukon River, and, subsequently, a less pronounced increase in isotope values before marine entry. Without this distinguishing feature, CWAK individuals from the Yukon (i.e. Figure 3b, column 4) are more likely to be misclassified with Kuskokwim or Nushagak fish, resulting in lower overall accuracy. Nonetheless, even applying the most conservative methods ( 90% confidence threshold on the restricted dataset ) we achieved 78% classification accuracy across all three datasets.

In addition to the ability to distinguish among major watersheds, the observed difference in isotope LA patterns between CWAK and non-CWAK individuals from the Yukon river suggests that the approaches described here may also be capable of resolving spatial structure within a single watershed. For example, genetics-based data in this dataset can produce known sub-watershed labels, which, given sufficient sample size, could be used to train a model to distinguish among genetic regions in the Yukon river-basin using the workflow described here. Previous work has demonstrated that integrating data sources such as genetics can enhance isotope-based assignment precision in some cases (Makhlouf et al., 2025). However, these preliminary observations suggest that similar approaches to the one outlined here may have the potential to improve assignment precision to the sub-basin scale without the need for additional data sources.

### Implications for Stock Assessment and Management

A tool to estimate provenance of marine-caught Chinook salmon in western Alaska has the potential to expand assessment of hypotheses of declining population health by allowing for an explicit assessment of marine mortality on stock specific health. The workflow developed here illustrates an approach that can be used alongside otolith LA-ICPMS data to provide probabilistic estimates of provenance and calibrated may be immediately useful for classifying adult Chinook salmon from western Alaska watersheds. Our dataset is unusually large for its type and includes samples collected across several years and from nearly all parts of the sampled watersheds. As such, we expect it to be broadly representative of the natural variability among years, and that the model presented here will generalize well to new years of data. However, due to sample size constraints we did not include a true validation set from a year excluded from the training and testing data, leaving open the possibility that some degree of overfitting to this dataset could affect assignment accuracy for unseen years. During exploratory analyses, we conducted a preliminary cross-year validation in which one year of Yukon data was withheld in each run and used as a validation set. Classification accuracy across years was generally consistent with testing performance, except for 2019, which performed moderately worse, possibly due to differences in sample preparation or data quality. While these preliminary results were not included in the final analysis they support the potential for this model to generalize to new years of data, which we plan to further evaluate as additional, currently unprocessed years of otolith data become available. Additionally, sample preparation procedures and LA-ICPMS specifications may vary among research groups, which could complicate direct comparisons between our dataset and others. This consideration also applies to analyses of juvenile Chinook salmon, which should display comparable isotopic patterns given that our model used only the freshwater region of the otolith. Nevertheless, differences in analytical approaches, particularly when working with much smaller otoliths, could influence results, and this assumption should be evaluated before large-scale application. Despite these caveats, we expect the patterns identified here to be robust across similar data types, particularly when paired with advanced time-series methods (ex. dynamic time warping (Arai et al., 2025 (*in review*)), or deep learning approaches which can accommodate variation in time-series length and minimize the need for preprocessing steps which may warp timeseries.

In addition to providing accurate watershed assignment estimates, the workflow presented here provides probability values which associate class estimates with a true measure of model uncertainty. In turn, managers or decision makers may use the result of the models presented here to make decisions based on flexible thresholds of misassignment risk which more accurately consider model uncertainty. In addition, these values can be appropriately used in probabilistic frameworks for assignment or structured decision making, thereby allowing more effective quantification of uncertainty through to management outcomes.

Despite an advancement in the ability to discriminate among rivers for western Alaska Chinook salmon, the workflow presented here remains limited for in-season management or rapid research applications. Otolith LA-ICPMS is costly and labor-intensive, making large-scale, real-time assessment of river origin for marine-caught samples impractical. Moreover, otolith collection is lethal, which may be prohibitive given the increasingly endangered status of Chinook salmon in this region. The model also relies on a large existing dataset with known river-of-origin labels derived from unique sampling efforts, a resource not available for most other species of concern. For example, comparable datasets do not exist for chum salmon, constraining the application of these methods to other similar species of concern in the region. However, if these data and logistical limitations are satisfied, the overall workflow is broadly generalizable and can be implemented to address similar challenges in other systems or for other species.

Machine learning methods provide an opportunity to uncover new insights in ecological datasets. In some cases, this may be critical for ecosystem-based management attempting to maintain fishing opportunity while protecting species and populations of conservation concern. In this case, enabling river-basin scale determination of provenance for marine-caught Chinook salmon allows managers to connect region-wide salmon population patterns with river-specific population health. Doing so may allow researchers to better understand how ecosystem processes at several spatial scales and across ecosystems are interconnected to informing overall ecosystem health. Developing this knowledge of ecosystem function across scales is essential for informing management and conservation strategies that sustain the services provided by healthy, functioning ecosystems.

## Acknowledgements

This work was not possible without the efforts of Dr. Sean Brennan, who motivated much of the data collection used in this study, and sought to advance isotope-based methods on fish otoliths for ecological research in Alaska. We thank the University of Washington’s eScience Institute Data Science and AI Accelerator program, which provided significant mentorship and resources to advance this project. We thank the efforts of Alaska Department of Fish and Game, and the Orutsararmiut Native Council for sample collection, and for the employees of the Alaska Department of Fish and Game Gene Conservation Lab who provided the genetics data used in this study. This work was supported by the Kuskokwim River Intertribal Fish Commission (KRITFC) and the Arctic-Yukon-Kuskokwim Sustainable Salmon Initiative (AYK-SSI).

## Author Contributions

Ben Makhlouf performed data analysis and wrote the manuscript. Valentina Staneva provided technical expertise and provided extensive consultation on project direction. Daniel E. Schindler directed the project and analysis, secured funding, and helped develop the manuscript. All authors contributed to editing the final manuscript.

