## Supplemental Figure 1 for "Machine learning applied to otolith microchemical data to discriminate stock of origin in salmon"

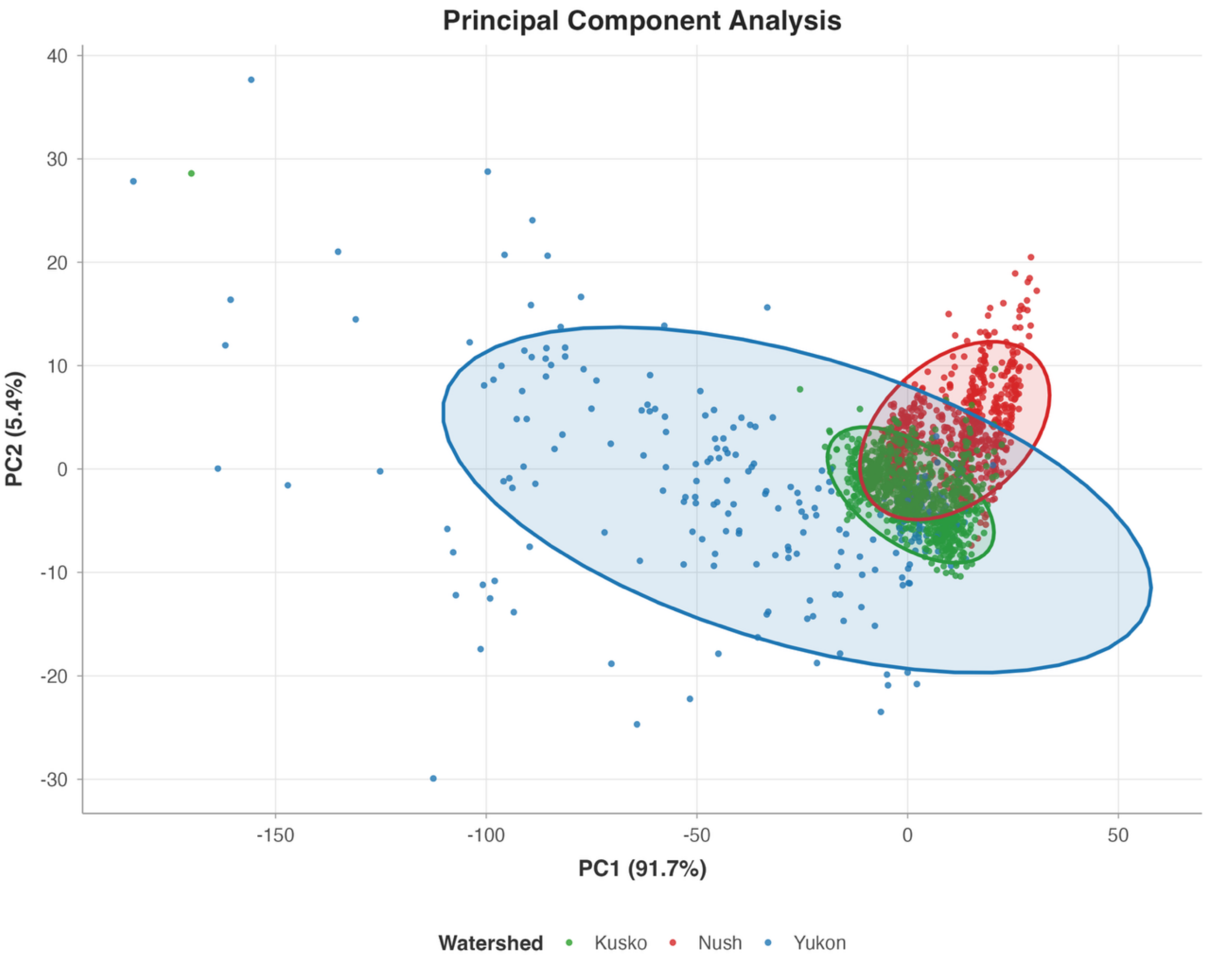


**Supp Figure 1:** PCA of GAM smoothed data with all Middle and Upper Yukon individuals removed, leaving only CWAK individuals. Data show a similar pattern to the PCA which included all individuals, but with many less individuals in the extreme range of both PC1 and PC2. This pattern illustrates the unique shape of the timeseries for many non-CWAK individuals compared to those in the lower watershed, which are more easily distinguished.
