## Supplemental Figure 2 for "Machine learning applied to otolith microchemical data to discriminate stock of origin in salmon"

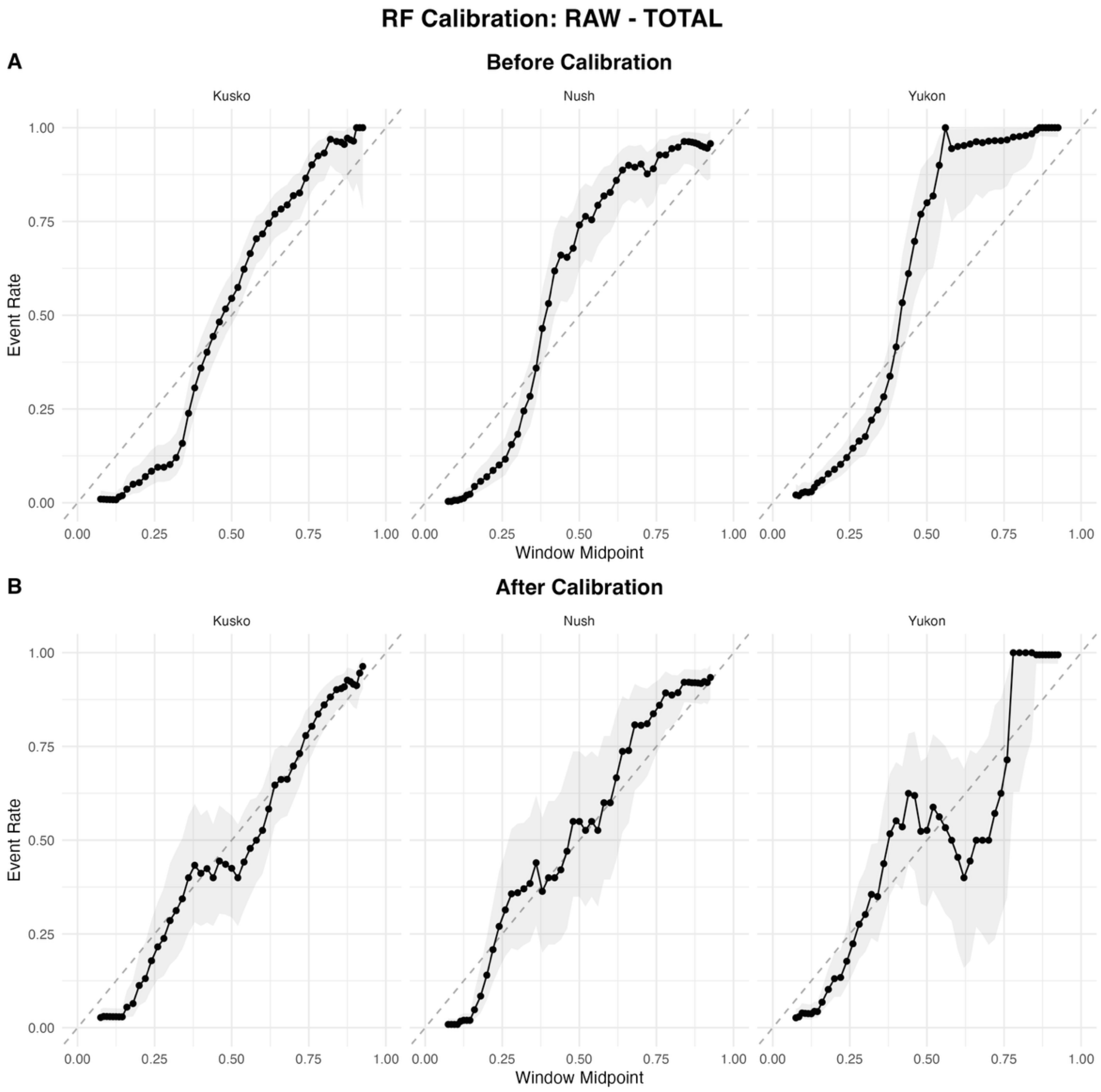


**Supp Figure 2:** Example diagrams showing the relationship between predicted probability value on the x axis and true class likelihood based on event rate on the y axis for random forest prediction on the total testing set. Predicted probability scores from the model (panel A) show a relatively linear relationship, indicating that probability scores are relatively well aligned before calibration. Applying a multinomial isotonic regression recalibrates probability scores to better match true class likelihood (panel B) , which is further confirmed by an increase in the brier score
